# Rapid and scalable characterization of CRISPR technologies using an *E. coli* cell-free transcription-translation system

**DOI:** 10.1101/169441

**Authors:** Ryan Marshall, Colin S. Maxwell, Scott P. Collins, Michelle L. Luo, Thomas Jacobsen, Chase L. Beisel, Vincent Noireaux

## Abstract

CRISPR-Cas systems have offered versatile technologies for genome engineering, yet their implementation has been outpaced by the ongoing discovery of new Cas nucleases and anti-CRISPR proteins. Here, we present the use of *E. coli* cell-free transcription-translation systems (TXTL) to vastly improve the speed and scalability of CRISPR characterization and validation. Unlike prior approaches that require protein purification or live cells, TXTL can express active CRISPR machinery from added plasmids and linear DNA, and TXTL can output quantitative dynamics of DNA cleavage and gene repression. To demonstrate the applicability of TXTL, we rapidly measure guide RNA-dependent DNA cleavage and gene repression for single- and multi-effector CRISPR-Cas systems, accurately predict the strength of gene repression in *E. coli*, quantify the inhibitory activity of anti-CRISPR proteins, and develop a fast and scalable high-throughput screen for protospacer-adjacent motifs. These examples underscore the potential of TXTL to facilitate the characterization and application of CRISPR technologies across their many uses.

## INTRODUCTION

CRISPR technologies have proven to be broadly useful genome-editing tools for biomolecular research, biotechnology, human health, and agriculture (Barrangou and Doudna, 2016). These technologies rely on RNA-guided nucleases derived from prokaryotic CRISPR-Cas immune systems (Mohanraju et al., 2016). The nuclease specifically cleaves DNA or RNA sequences complementary to the guide portion of the RNA and flanked by a protospacer-adjacent motif (PAM), allowing programmable DNA damage or repair-mediated genome editing. Furthermore, by disrupting endonuclease activity, these nucleases can be readily converted into programmable nucleic-acid binding proteins, and used for applications in gene regulation, base editing, and real-time imaging (Chen et al., 2016; Dominguez et al., 2016; Komor et al., 2017; Nelles et al., 2016).

While the vast majority of CRISPR-based technologies have relied on the DNA-targeting Cas9 nuclease from *Streptococcus pyogenes* (Barrangou and Doudna, 2016; Jinek et al., 2012), nature boasts a diverse collection of CRISPR-Cas systems that are currently sub-divided into two classes, six types, and 33 subtypes (Koonin et al., 2017; Makarova et al., 2015; Shmakov et al., 2015). Interrogation of the emerging subtypes have revealed nucleases with widely varying properties that can be smaller, recognize different PAMs, degrade DNA, target RNA, exhibit reduced propensity for off-target effects, or are more amenable to multiplexing in comparison to SpyCas9 (Abudayyeh et al., 2016; Kim et al., 2017; Kleinstiver et al., 2016; Mulepati and Bailey, 2013; Zetsche et al., 2017). Separately, the discovery of anti-CRISPR proteins that inhibit Type I-E, I-F, II-A, and II-C CRISPR-Cas systems offer means to tightly control nuclease activity (Bondy-Denomy et al., 2015; Pawluk et al., 2016a; Pawluk et al., 2016b; Rauch et al., 2017), with the likely existence of similar inhibitors for the other subtypes. However, despite this diversity, the vast majority of these proteins have been slow to be adopted as CRISPR technologies.

One major bottleneck is the characterization of these proteins’ basic properties and functions. To date, characterization has been performed with multiple methodologies based on *in vitro* biochemical assays or live cells. More recently, these assays have been modified for high-throughput analysis of PAM binding requirements, off-target propensities, or large libraries of guide RNAs through next-generation sequencing or imaging of arrayed nucleotides (Abudayyeh et al., 2016; Boyle et al., 2017; Jiang et al., 2013; Karvelis et al., 2015; Kleinstiver et al., 2016; Leenay et al., 2016; Pattanayak et al., 2013; Tsai et al., 2017). However, these assays consistently require days to weeks to perform due to the requirement for protein purification or for culturing and transforming live cells. Furthermore, these assays scale poorly when testing large sets of proteins or guide RNAs. Given the growing abundance of known CRISPR nucleases, the ease in guide rna design, and the growing prevalence of factors factors that interface with CRISPR-Cas systems, there remains a pressing need to develop rapid and scalable characterization methodologies.

Here, we address this need using an *Escherichia coli* cell-free transcription-translation system (TXTL) (Garamella et al., 2016; Shin and Noireaux, 2012). TXTL recapitulates gene expression and enzymatic activity following the addition of template DNA, allowing the quantitative and dynamic measurement of nuclease binding and cleavage at microliter scale in a few hours -- all without protein purification or live cells. We demonstrate the utility of TXTL across a diverse set of Cas nucleases and show how it can be used to predict guide RNA activity *in vivo*, characterize anti-CRISPR proteins, and elucidate recognized PAMs. Based on these findings, we expect TXTL to provide a powerful characterization tool for the expanding universe of CRISPR-Cas systems and proteins, and we show the applicability of TXTL beyond biomanufacturing, diagnostics, and genetic circuit prototyping (Carlson et al., 2011; Dudley et al., 2014; Garamella et al., 2016; Gootenberg et al., 2017; Hockenberry and Jewett, 2012; Jewett et al., 2008; Kanter et al., 2007; Karzbrun et al., 2014; Pardee et al., 2016; Sun et al., 2014; Swartz, 2006; Takahashi et al., 2015a; Takahashi et al., 2015b; Tayar A., 2015).

## RESULTS

### SpyCas9 and dSpyCas9 exhibit robust activity in TXTL

We initially examined the activity of the SpyCas9 nuclease. To monitor the dynamics of DNA cleavage by SpyCas9, we designed single-guide RNAs (sgRNAs) that target within the promoter, 5’ untranslated region (UTR), and coding sequence of a deGFP reporter construct, a slightly modified version of eGFP with identical fluorescent properties (Shin and Noireaux, 2012). We then measured the dynamics of deGFP production following the addition of the SpyCas9 plasmid, linear DNA encoding the sgRNA, and the deGFP reporter plasmid (Figure 1A). The resulting fluorescence values were converted to a true reporter concentration based on a calibration curve made with purified eGFP. We found that the three tested sgRNAs resulted in greatly reduced deGFP concentrations in comparison to a non-targeting sgRNA after 16 hours in the TXTL reaction (Figure 1B). Measurable repression was observed beginning after less than one hour into the TXTL reaction, indicating that this time span was required to express and assemble the Cas9-sgRNA ribonucleoprotein complex (RNP) and for the complex to bind and cleave the target DNA (Figure S1A,B). The rate of deGFP production then dropped quickly to approximately zero, consistent with irreversible DNA cleavage as confirmed by PCR amplification of the target site (Figure S1C). Interestingly, the onset and of deGFP repression and the rate at which deGFP production dropped varied depending on the sgRNA used (Figure S1A,B), suggesting that varying efficiency by which different guides targeted Cas9 to DNA. These results indicate that an active SpyCas9-sgRNA complex can be expressed directly in TXTL, providing a dynamic and quantitative readout of nuclease activity in a few hours.

**Figure 1.**
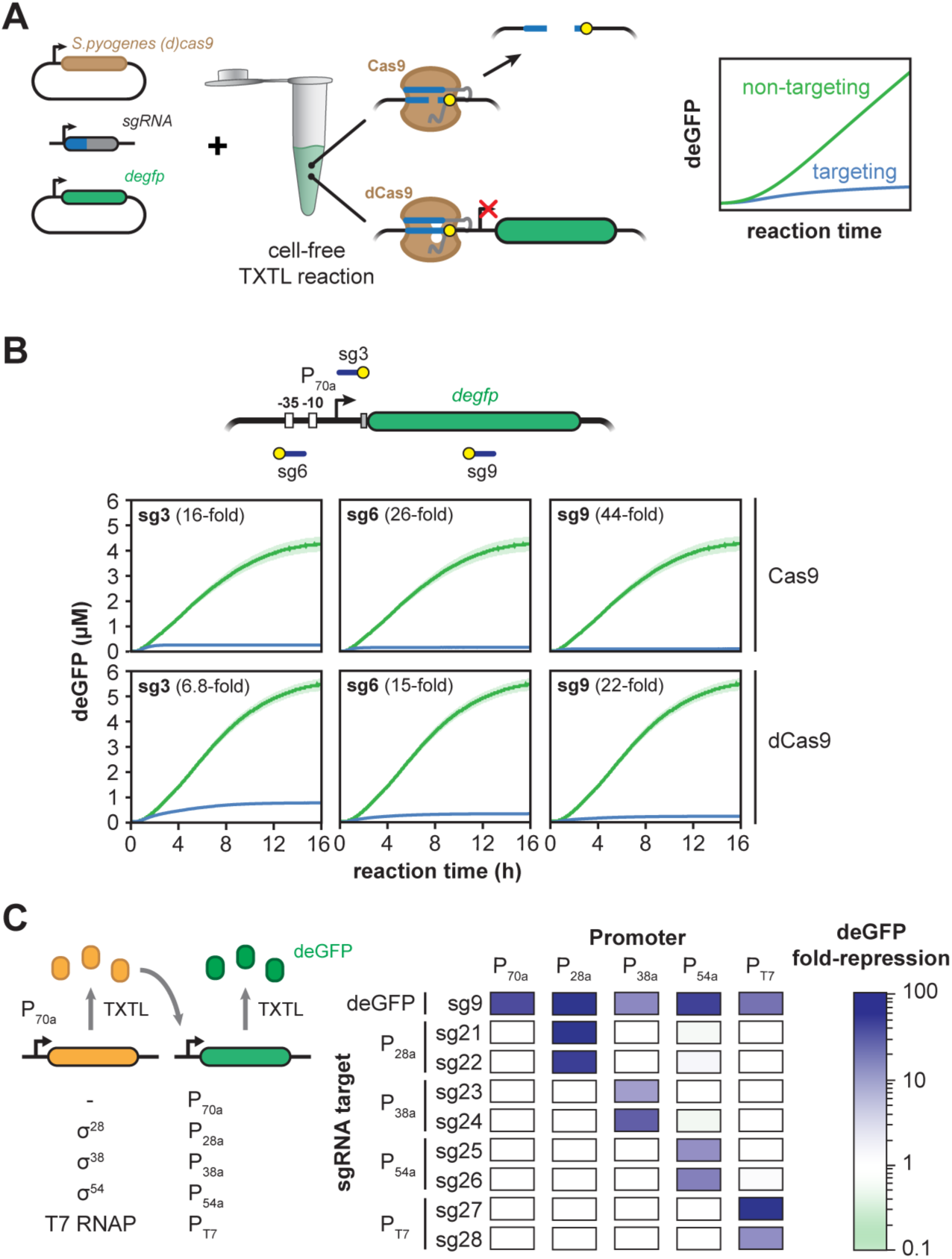
*S. pyogenes* Cas9 functions efficiently in TXTL. **A.** Schematic of using TXTL to dynamically and quantitatively measure the activity of Cas9 and the catalytically-dead Cas9 (dCas9) repression based on repressing expression of a deGFP reporter construct. Plasmids expressing a reporter gene (deGFP) and SpyCas9 or dSpyCas9 as well as linear DNA expressing an sgRNA are added to the cell-free reaction. sgRNAs direct the Cas9 or dCas9 ribonucleoprotein (RNP) complex to the reporter construct, leading to cleavage or binding by the RNP, respectively. Cleavage or binding can be monitored based on GFP fluorescence over time in a microplate reader. **B.** Time series showing deGFP concentration for cell-free reactions expressing either Cas9 or dCas9 and a non-targeting sgRNA (green) or targeting sgRNAs (blue). Target locations include the sequence matching the guide (blue line) and the PAM (yellow circle). **C.** Alternative sigma factors σ28, σ38, and σ54 and the T7 polymerase can be expressed in TXTL from the P_70a_ promoter and activate their cognate promoters P_28a_, P_38a_, P_54a_, and P_T7_, respectively. A matrix showing dSpyCas9-based repression of promoters dependent on σ28, σ38, σ54 and the T7 polymerase is shown. An sgRNA targeting each promoter or the GFP gene body was expressed along with each sigma factor or polymerase and a reporter gene driven by the sigma factor of its cognate promoter. For each condition, repression was calculated after a reaction time of 16 hours in comparison to a non-targeting sgRNA.

We also interrogated the activity of the catalytically-dead version of SpyCas9 (dSpyCas9) commonly used for programmable DNA binding and gene regulation -- also called CRISPRi and CRISPRa (Gilbert et al., 2014; Qi et al., 2013). DNA binding by dSpyCas9 can block transcriptional initiation or elongation of RNA polymerase in bacteria, offering a means to link DNA binding with reporter expression in TXTL (Bikard et al., 2013; Qi et al., 2013). To assess the expression and regulatory activity of dSpyCas9, we measured the concentration of deGFP over time in TXTL reactions for the same three targeting sgRNAs and the non-targeting sgRNA. Similar to reactions expressing SpyCas9, reactions expressing dSpyCas9 exhibited consistent deGFP repression, and the rate of deGFP production dropped after less an hour (Figure S1A,B). The catalytically-dead version repressed expression less strongly and exhibited some deGFP production even at the end of the TXTL reaction (Figures 1B, S1A). We confirmed that the dSpyCas9 did not cleave DNA by PCR amplification across the target location (Figure S1B). We also observed different extents of deGFP fold-repression across the three targeting sgRNAs similar to repression strengths reported in bacteria (Bikard et al., 2013; Qi et al., 2013). These results illustrate that dSpyCas9 functions efficiently to repress gene expression in TXTL. Given the larger span of time in which regulatory activity can be observed for dSpyCas9 than for SpyCas9 (Figure S1A), we relied on CRISPR-based gene repression for all subsequent analyses.

To assess the scalability of TXTL reactions, we expanded from one promoter to four promoters each targeted by two sgRNAs and driving expression of deGFP. Each promoter is dependent on a unique alternative sigma factor (σ^28^, σ^38^, σ^54^) or the T7 RNA polymerase, where each transcription factor is supplied on an added expression plasmid under the control of a σ^70^ promoter (Garamella et al., 2016; Shin and Noireaux, 2012) (Figure 1C). By including a deGFP-targeting sgRNA, a non-targeting sgRNA, and a σ^70^ promoter, we tested a total of 50 promoter-sgRNA combinations, with each combination tested in triplicate. The TXTL reactions confirmed that measurable repression was only observed when an sgRNA was matched with its target (Figure 1C), with the strength of repression ranging between 7-fold and 105-fold. We also targeted binding sites for the NtrC operator sites within the σ^54^ promoter (Figure S2), demonstrating that these sites were important for reporter expression. In total, we showed that TXTL can be used to rapidly and scalably assess sgRNA activity.

### Multiple factors impact the measured activity of dSpyCas9 in TXTL

After characterizing SpyCas9 and dSpyCas9 in TXTL, we sought to determine parameters that affect the measured activity. First, we found that varying the amount of the dSpyCas9 plasmid reduced the total amount of GFP produced in a TXTL reaction (Figure S2A), suggesting that dSpyCas9 levels were limiting under these reaction conditions. Surprisingly, adding more dSpyCas9 plasmid elevated deGFP production for the non-targeting sgRNA, underscoring the need to add the same total amount of DNA when performing comparative studies such as when comparing the activity of different targeting sgRNAs.

Second, we evaluated the impact of destabilizing deGFP to emulate reporter turnover or dilution *in vivo.* We appended an ssrA degron tag recognized by the ClpXP protease to the C-terminus of deGFP (Figure 2B) (Schmidt et al., 2009). Appending the ssrA tag to deGFP resulted in an apparent enhancement of dSpyCas9-based repression (Figure 2B) due to the inability of deGFP to accumulate in the TXTL reaction (Figure S2B). This apparent enhancement offers a means to measure CRISPR-based repression under conditions that mimic dilution *in vivo.*

**Figure 2.**
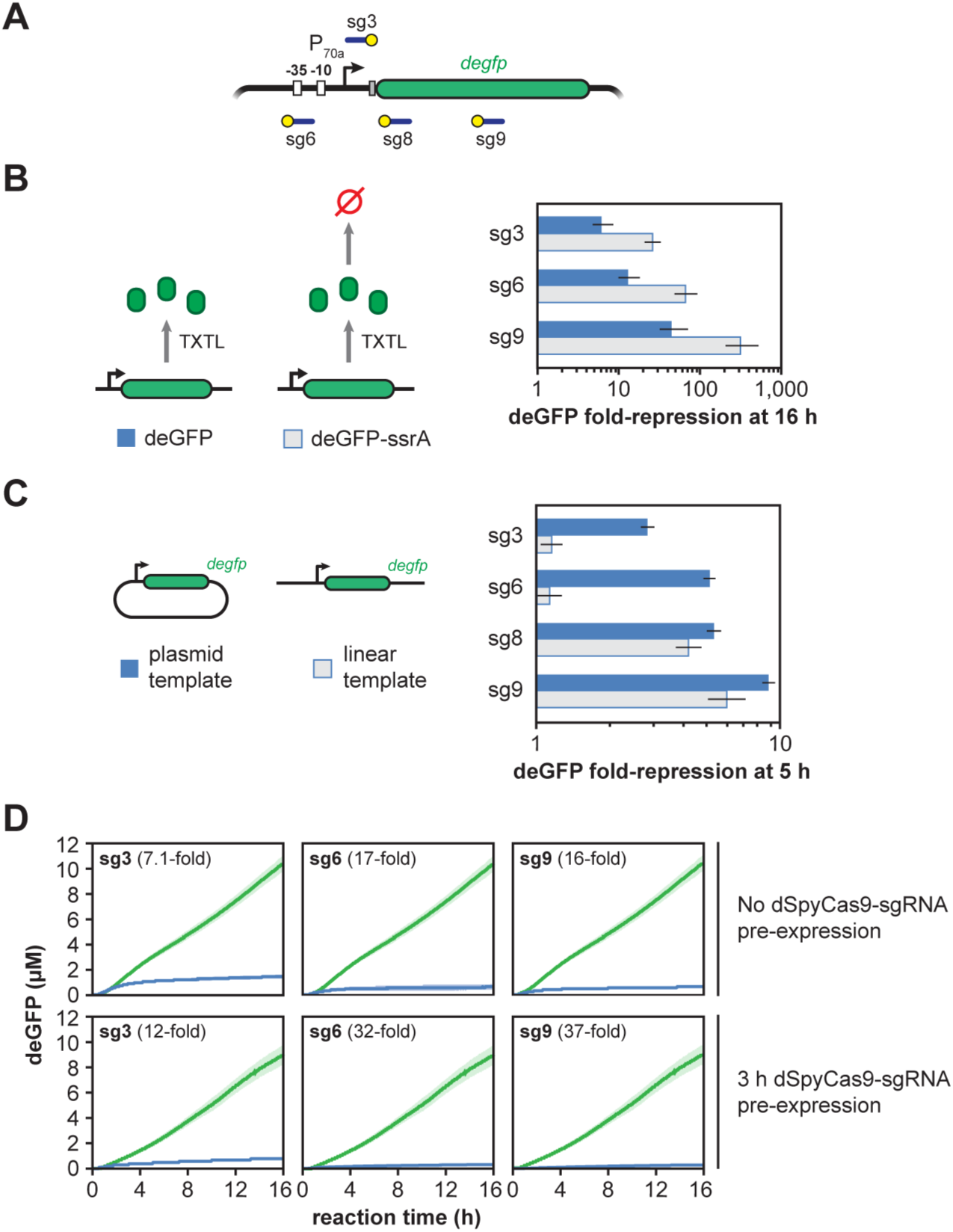
Multiple factors affect dSpyCas9-based repression of reporter gene expression in TXTL. **A.** Locations within the deGFP reporter construct targeted with the sgRNAs. Target locations include the sequence matching the guide (blue line) and the PAM (yellow circle). **B.** Fold-repression for reporter constructs encoding deGFP or deGFP-ssrA. The ssrA degron tag is recognized by the ClpXP protease that results in rapid turnover of the fusion protein. Fold-repression is the ratio of deGFP concentrations after 16 hours of reaction for the non-targeting (green) over the targeting (blue) sgRNA. **C.** Fold-repression produced by a TXTL reaction when deGFP is expressed from either a targeted linear or plasmid construct. **D.** Time series showing deGFP concentration in TXTL for cell-free reactions expressing dSpyCas9 and a targeting sgRNA. The reporter plasmid is added to the reaction either at the same time as dCas9 and the sgRNA (top row) or after a 3 hour pre-expression of dSpyCas9 and the sgRNA (bottom row).

We next evaluated the impact of targeting linear versus plasmid DNA. Linear DNA can be generated without cloning and thus can be readily used in TXTL (Marshall et al., 2017; Sun et al., 2014). However, some Cas nucleases have been shown to require supercoiled DNA for binding (Westra et al., 2012), suggesting that relaxed DNA may not efficiently recognize targets encoded on linear DNA. We measured the extent of dSpyCas9-based repression when targeting a plasmid or linear version of the deGFP reporter construct. The linear DNA was generated by PCR including flanks over one kilobase away from the closest target site, mitigating potential effects from the end of the dsDNA molecule. We measured the fold-repression of the deGFP produced by the reaction after five hours, which is shorter than our previous experiments because linear templates degrade after that time in our TXTL system (Marshall et al., 2017). We found that dSpyCas9 was capable of repressing gene expression when targeting the transcribed sequence of the reporter gene on the linear template, but not when targeting the promoter (Figure 2C). These results suggest that the dSpyCas9 can block elongation but not initiation of the *E. coli* RNA polymerase on linear DNA. They also indicate that Cas nucleases interact differently with linear targets and plasmid targets, underscoring the need for further interrogation.

When DNA expressing dSpyCas9, sgRNA, and reporter plasmid DNA were added together at the beginning of a TXTL experiment, we observed a transient period in which the reporter gene was expressed before the onset of dSpyCas9-based repression (Figure 1B). We hypothesized that this period of transient expression was due to the slow assembly speed of the dSpyCas9-sgRNA RNP complex. To test this hypothesis, we varied our initial protocol to pre-express dSpyCas9 and the sgRNA before adding the reporter plasmid for three hours. We reasoned that this period of pre-expression would allow the RNP to assemble and that this would shorten the time until we observed repression of the reporter gene. Consistent with our hypothesis, pre-expressing dSpyCas9 and the sgRNA reduced the time before the reporter gene was repressed (Figure 2C), with measurable repression occurring as fast as six minutes, depending on the sgRNA used (Figure S2C,D). We also evaluated the effect of pre-expressing dSpyCas9 in cells prior to generating the TXTL lysate (Figure S2E). Interestingly, pre-expression of dSpyCas9 generally did not enhance repression (Figure S2F), suggesting that expression of the sgRNA and assembly of the ribonucleoprotein complex strongly contribute to the delayed onset of dSpyCas9-based repression. Taken together, these results suggest that dCas9 RNP assembly is relatively slow (on the order of 30 minutes), but that DNA binding is faster (on the order of 5 minutes).

### The strength of dSpyCas9-based repression strongly correlates between TXTL and *E. coli.*

Given the speed and scalability of employing TXTL to characterize CRISPR nucleases and guide RNAs, an ensuing question is how well the quantified activity correlates to *in vivo* settings and recapitulates known phenomena (Chappell et al., 2013). To begin addressing this question, we opted to compare dSpyCas9-based repression in TXTL and in *E. coli* by targeting 19 different locations within the deGFP reporter plasmid (Figure 3). The particular locations were chosen to also evaluate the impact of strong PAMs (NGG) and weak PAMs (NAG) as well as targeting the template or non-template strand within the promoter or transcribed region. We chose these comparisons because of poor recognition of NAG PAMs than NGG PAMs by dCas9 (Boyle et al., 2017; Jiang et al., 2013) and reduced gene repression in bacteria when targeting the template strand within the transcribed region versus any other location (Bikard et al., 2013; Qi et al., 2013). The experiments were conducted by encoding the dSpyCas9, sgRNA, and deGFP reporter on separate, compatible plasmids for parallel testing in TXTL and in *E. coli*, and fold-repression was calculated based on endpoint measurements of deGFP in comparison to a non-targeting sgRNA control.

**Figure 3.**
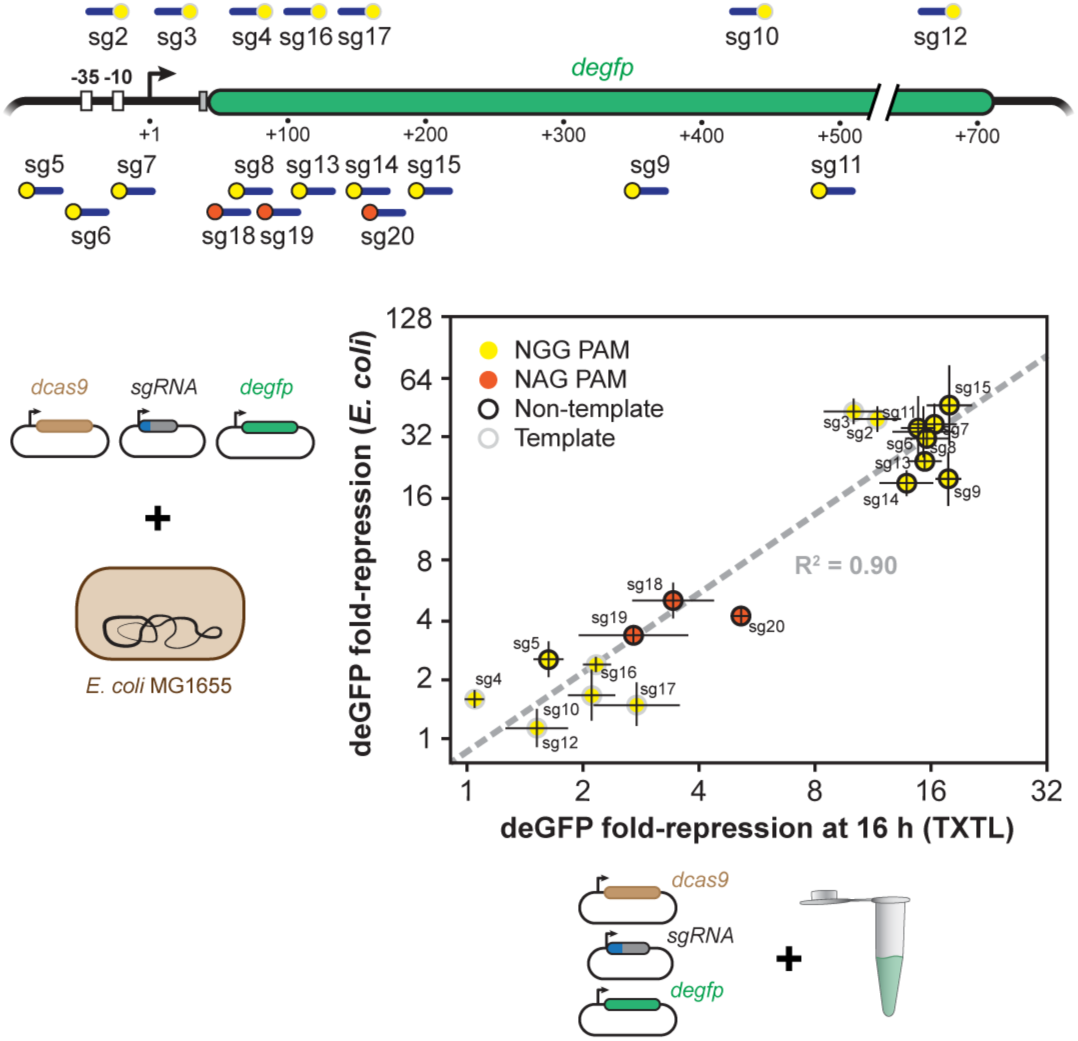
dSpyCas9-based repression *in vivo* and *in vitro* are well-correlated. A schematic of where each guide binds in the GFP promoter and gene body is shown (top). The location of the target and PAM are indicated by a blue line and a yellow or orange dot, respectively. The fold-repression of GFP production by CRISPRi for a guide RNA *in vivo* and *in vitro* is shown (bottom). Points are colored by whether the guide is adjacent to an NGG (yellow) or NAG (orange) PAM, and whether the sgRNA targets the non-template (black ring) or non-template (gray ring) strand.

These experiments revealed a strong correlation (R^2^ = 0.90) between the measurements in TXTL and in *E. coli* (Figure 3). Consistent with this correlation, we observed greatly reduced repression for targets flanked by NAG PAMs versus NGG PAMs. Furthermore, repression was consistently weaker when targeting the template strand of the transcribed region in comparison to targeting the non-template strand of the transcribed region or either strand within the vicinity of the promoter. This direct comparison therefore showed that TXTL can reasonably predict *in vivo* activity of CRISPR-Cas systems, at least for gene repression by dSpyCas9 in *E. coli.*

### The activity of other CRISPR nucleases can be measured with TXTL

Our results showing that active SpyCas9 RNPs can be expressed in TXTL suggested that other CRISPR-Cas systems would also function in TXTL. To test this, we evaluated effector proteins from two other CRISPR-Cas systems: the single-effector nuclease Cpf1 from the Type V-A system in *Francisella novicida*, and the multi-subunit complex Cascade from the Type I-E system in *E. coli* (Figure 4). These were selected because Cpf1 nucleases exhibits many desirable properties over Cas9 nucleases while Type I CRISPR-Cas systems are the most prevalent system type in nature but are more challenging to characterize because multiple proteins form the effector complex (Makarova et al., 2015; Zetsche et al., 2015). Previous studies have shown that both systems are both capable of programmable gene repression in bacteria when using a catalytically-dead version of the nuclease (dFnCpf1) or by expressing Cascade in the absence of the Cas3 endonuclease, respectively (Leenay et al., 2016; Luo et al., 2015; Rath et al., 2015; Zetsche et al., 2015). To test CRISPR-based repression for each system, we designed three guide RNAs (a mature guide RNA for Cpf1, a repeat-spacer-repeat for Cascade) targeting distinct sites within the P70a promoter, with each site flanked by an appropriate PAM.

**Figure 4.**
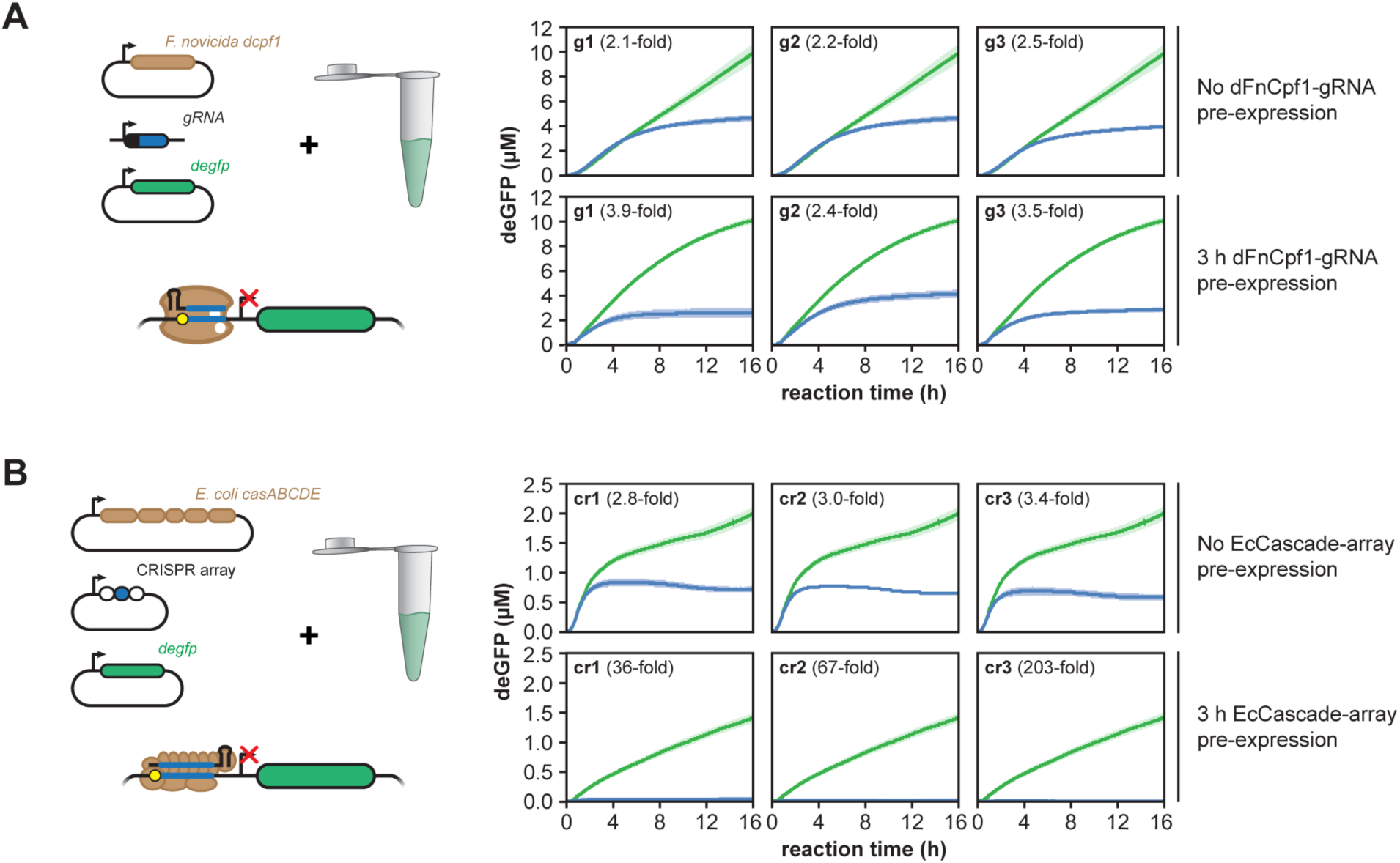
Single effector and multi-protein effector Cas proteins function efficiently in TXTL. Time series of reporter gene expression in TXTL for cell-free reactions expressing **A.** a catalytically inactive version of the Type V-A Cpf1 nuclease from *Francisella novicida* or **B.** the Type I-E Cascade complex from *E. coli.* The protein or set of proteins were expressed along with a non-targeting guide RNA (green) or one of three guide RNAs (blue) designed to target the promoter of the deGFP reporter construct. The reporter plasmid is added to the reaction either at the same time as the constructs expressing the Cas protein(s) and the guide RNA (top row) or after three hours of pre-expression (bottom row). The guide RNAs were expressed as a mature CRISPR RNA (FnCpf1) or as a repeat-spacer-repeat (EcCascade).

We first found that dFnCpf1 was capable of repressing gene expression with each of the three guide RNAs that targeted the reporter gene promoter (Figure 4A). The measured repression by dFnCpf1 after 16 hours of reaction was weaker than that achieved by dSpyCas9, which we attribute to the longer delay in the onset of repression (Figure S3A,C). Interestingly, pre-expressing dFnCpf1 and its guide RNA for three hours reduced the delay in measurable repression by approximately two hours. This finding suggests that complex assembly and DNA binding is slower for the FnCpf1-gRNA complex than for the dSpyCas9-sgRNA complex.

We also found that EcCascade could elicit gene repression in TXTL despite the need to coordinately express five proteins. The total reduction of GFP produced by the reaction was modest because of the delay in the onset of strong repression (Figures 4B and S3B,D). However, pre-expressing EcCascade and the CRISPR RNA strongly reduced the total amount of GFP produced due to strong repression shortly after the addition of the reporter plasmid (Figures 4B and S3B,D), indicating that the complex -- once expressed and assembled -- rapidly binds DNA and efficiently blocks RNA polymerase recruitment. Unexpectedly, deGFP production was consistently lower when expressing EcCascade versus any of the other effector proteins that was exacerbated when EcCascade was pre-expressed (Figure 4B), suggesting that EcCascade is interfering with the expression or activity in deGFP. In total, we show that TXTL can be extended to the characterization of CRISPR-Cas systems requiring both single-effector and multi-protein effector complexes.

### TXTL can quantify the inhibitory activity of anti-CRISPR proteins

Recently, the discovery of anti-CRISPR proteins that bind and inhibit Cascade and Cas3 from Type I-E and Type I-F CRISPR-Cas systems and Cas9 from Type II-A and II-C systems raised the potential of using these proteins to tightly regulate genome editing and gene regulation (Bondy-Denomy et al., 2015; Pawluk et al., 2016a; Pawluk et al., 2016b; Rauch et al., 2017). We asked if TXTL could be used to rapidly assess the inhibitory activity of potential anti-CRISPR proteins. We focused on AcrIIA2 and AcrIIA4, two anti-CRISPR proteins that were recently reported to inhibit the activity of SpyCas9 *in vitro* and in human cells (Rauch et al., 2017). To measure the inhibitory activity of AcrIIA2 and AcrIIA4 against SpyCas9 in TXTL, we encoded each protein on a linear expression construct that was expressed for two hours in the lysate prepared from cells expressing dSpyCas9 (Figures 5A, S2E). We then added DNA encoding the GFP reporter plasmid and linear DNA encoding a targeting or non-targeting sgRNA and measured deGFP fluorescence over time.

**Figure 5.**
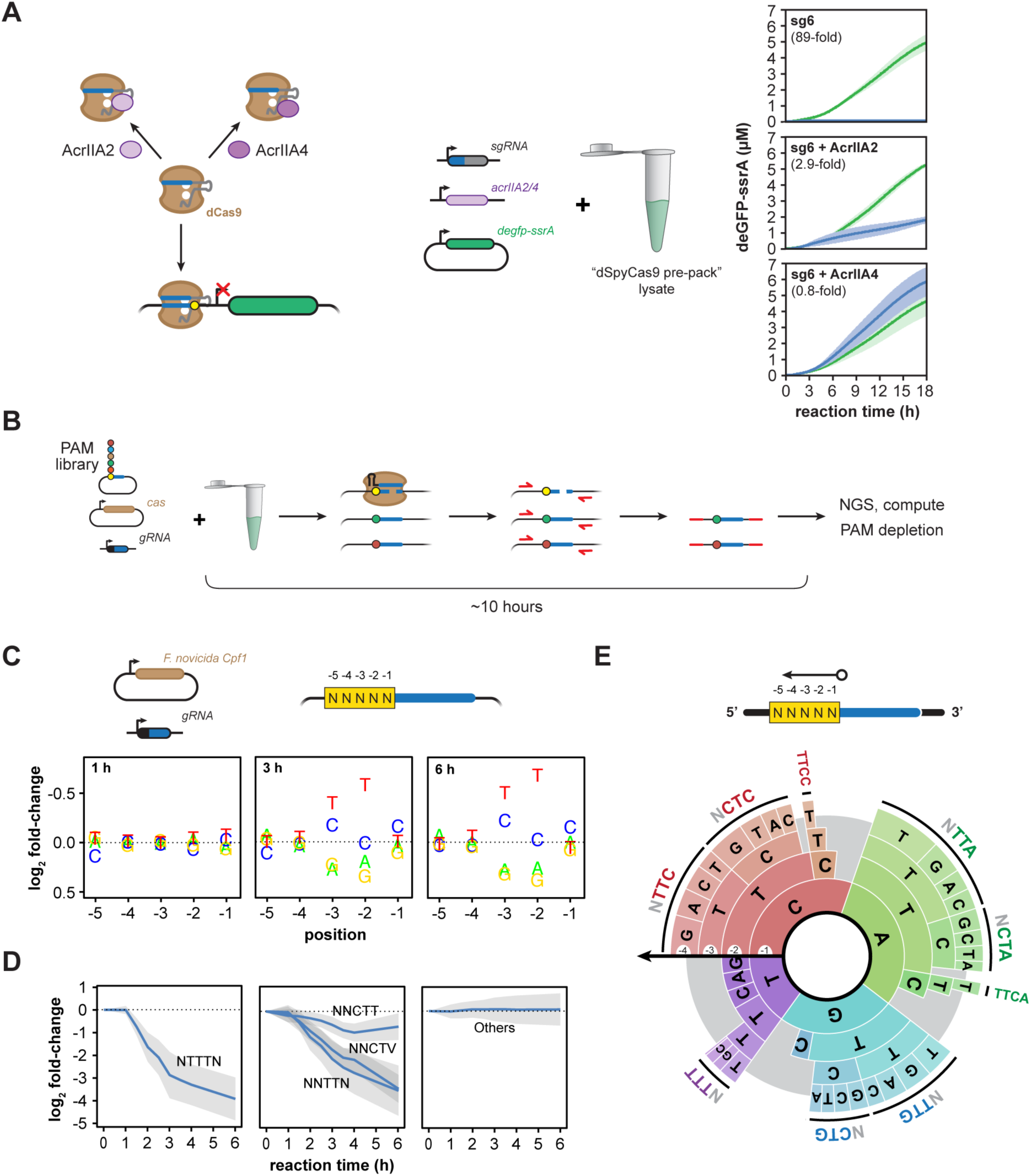
Using TXTL to characterize anti-CRISPR proteins and to determine PAMs. **A.** Time series of deGFP-ssrA expression in TXTL for cell-free reactions also expressing dSpyCas9, an sgRNA, and one of two anti-CRISPR proteins, AcrIIA2 and AcrIIA4, shown to inhibit SpyCas9 activity. Each reaction was performed with a targeting sgRNA (blue) or a non-targeting sgRNA (green). **B.** Schematic of a TXTL-based cleavage assay to determine the PAM sequences recognized by Cas nucleases. Plasmids expressing the Cas nuclease and a guide RNA targeting a sequence flanked by randomized base pairs are combined and incubated in TXTL. Plasmids containing valid PAMs are cleaved by the Cas nuclease, while plasmids lacking a PAM are uncut. Sequencing libraries are then created by PCR amplifying across the target site, resulting in depletion of PAM sequences in the subsequent sequencing library. **C.** Plots showing the fold-change in the representation of a nucleotide at each variable position in the PAM library in comparison to the original PAM library. Note that the y-axis is inverted to highlight nucleotides that are depleted. **D.** Time series showing the depletion of selected motifs matching the consensus sequence in the sequencing libraries are shown. Error bars show the standard deviation of the fold-change. **E.** A PAM wheel showing the determined PAM sequences recognized by FnCpf1. PAM sequences are read proceeding from the outside to the inside of the circle, and the arc length directly correlates with the extent of PAM depletion. The −5 position was not shown for clarity.

We found that both AcrIIA2 and AcrIIA4 counteracted gene repression by dSpyCas9. However, AcrIIA4 fully restored the amount of deGFP produced by the reaction compared to the non-targeting control, whereas AcrIIA2 only restored deGFP production by 34%. Therefore, TXTL can be used to quantify the inhibitory activity of anti-CRISPR proteins, where the resulting values can guide the rapid identification of potent anti-CRISPR proteins that can more effectively inhibit Cas nuclease activity.

### TXTL offers a rapid means of elucidating CRISPR PAMs

One of the major barriers to the functional characterization of new Cas nucleases is determining the recognized PAM sequences. While numerous experimental methods have been developed for PAM determination, they consistently rely on *in vitro* assays that require protein purification or on cell-based assays that require culturing and transforming of live cells (Karvelis et al., 2017; Leenay and Beisel, 2017). We therefore asked if TXTL could also be used as the basis of PAM determination assays to avoid protein purification and exploit the large library sizes that can be screened with *in vitro* assays. Paralleling prior *in vitro* and *in vivo* DNA cleavage assays (Jiang et al., 2013; Zetsche et al., 2015), our devised assay relies on introducing a library of potential PAM sequences flanking a site targeted by an expressed guide RNA and Cas nuclease (Figure 5B). After incubating the pooled PAM library and DNA encoding the Cas nuclease and guide RNA, the pool of uncleaved target sequences is PCR-amplified and subjected to next-generation sequencing. By comparing the relative frequency of individual sequences within the library before and after sequencing, we can quantify the depletion of each library member and therefore how well the nuclease recognizes each sequence as a PAM. Critically, the assay can be completed in ~10 hours from when the DNA constructs are in hand to when PCR products are submitted for sequencing.

As a proof-of-principle demonstration, we assessed the PAM requirements of the well-characterized FnCpf1 nuclease. Previous assays have revealed that FnCpf1 requires a TTN motif on the 5’ end of the target for efficient cleavage using the guide RNA-centric orientation, although CTN can also be recognized (Fonfara et al., 2016; Leenay et al., 2016; Zetsche et al., 2015). We created a five-nucleotide library of potential PAM sequences 5’ to the guide RNA target. *In vitro* PAM assays are known to yield less specific PAM sequences for higher nuclease concentrations (Karvelis et al., 2015), so we assessed the determined PAMs in our assay for reaction times ranging from one hour to six hours following the addition of the FnCpf1 and guide RNA expression constructs (Figure S4). Figure 5 depicts the identified PAM sequences based on the depletion of individual nucleotides at each position (Figure 5C) or specific motifs (Figure 5D) as well as a PAM wheel capturing the relative depletion of each sequence across the library (Figure 5E) (Leenay et al., 2016). We did not include the −5 position in the PAM wheel because this position showed no appreciable bias across the four possible nucleotides.

Our TXTL-based PAM assay recapitulated the canonical 5’ TTN PAM for FnCpf1 while revealing other distinct features. At a reaction time of six hours, NTTTN was the most active motif. Within this motif, ATTTA was the most depleted by ~3-fold more than the next most depleted sequences (Data file S1). We further found that NNCTN also supported efficient cleavage in line with previous reports (Fonfara et al., 2016; Zetsche et al., 2015), while a T at the −1 position was detrimental to cleavage. Taken together, these results indicate that a TXTL-based assay can elucidate PAM requirements for CRISPR-Cas systems, where the assay opens the door to the rapid and scalable characterization of PAM requirements for novel Cas nucleases.

## DISCUSSION

We have demonstrated that *E. coli* cell-free TXTL can be used for the rapid characterization of CRISPR nucleases, guide RNAs, and anti-CRISPR proteins. Unlike *in vitro* biochemical assays or cell-based assays, TXTL does not require any protein purification or cell culturing and transformation. TXTL also allows exquisite control over the reaction conditions and the amount of the DNA templates, and it can provide a dynamic and quantitative readout of nuclease activity within a few hours. We exploit these capabilities by (1) demonstrating DNA binding and cleavage by multiple types of Cas nucleases, (2) extracting the dynamics of component expression, complex formation, and DNA targeting, (3) show that the strength of gene repression strongly correlates between TXTL and *E. coli*, (4) characterize the inhibitory strength of anti-CRISPR proteins, and (5) devise a rapid PAM determination assay. With recent advances in DNA synthesis as well as liquid handling systems, these approaches could be readily scaled to hundreds of reaction conditions or constructs.

CRISPR-Cas systems are remarkably diverse, with significant sequence, structural, and functional diversity (Koonin et al., 2017). This diversity exists even within a single subtype; for instance, Cas9 proteins from the well-characterized Type II-A subtype can exhibit less than 10% sequence homology at the amino-acid level and show a range of recognized PAM lengths and sequences (Fonfara et al., 2014). However, only a few representative nucleases have been characterized for the other subtypes. This is particularly striking for Type I and III CRISPR-Cas systems, the most prevalent types in prokaryotes. The issue is that these systems rely on multiple proteins to form the effector complex, requiring the purification or expression of multiple proteins in defined stoichiometries that has complicated their widespread characterization. TXTL is ideally suited to characterize these multi-subunit effector complexes from these systems because linear, chemically synthesized DNA encoding each subunit can be combined in a single TXTL reaction. The expressed complex can then be characterized in a variety of ways, such as determining its assembly kinetics (see below) and PAM requirements. Evaluating numerous systems from one subtype could help reveal how CRISPR-Cas systems evolved and the structural basis of PAM recognition through mapping sequence-function relationships.

While our results demonstrate the promise of TXTL for characterizing CRISPR-Cas systems, there are multiple opportunities to further expand the utility of TXTL. For example, TXTL could be used to investigate spacer acquisition across the diversity of Cas acquisition proteins found in nature. Our demonstration of the strong correlation between CRISPR-based repression *in vivo* and *in vitro* also suggests that TXTL could provide a rapid means to functionally validate guide RNAs before they are deployed for genome editing. However, more research is need to determine how binding or cleavage activity in TXTL correlates to genome editing and gene regulation in prokaryotic or eukaryotic cells or whether cellular factors such as DNA repair pathways or heterochromatin formation principally determine targeting efficiency. Next, we envision the development of further high-throughput screening assays using TXTL, such as assessing the sequence-dependence of guide RNA activity. Finally, measuring the time to repression with or without pre-expression of the CRISPR machinery opens the potential of using TXTL to rapidly measure the kinetics of ribonucleoprotein complex assembly and function under *in vivo*-like conditions. Through the demonstrations reported here and through further extensions, TXTL has the potential to make a widespread impact on the characterization of CRISPR-Cas systems and their transition into a new generation of CRISPR technologies.

## ACKNOWLEDGEMENTS

We thank Ryan Leenay for help generating the FnCpf1 PAM wheel, and Jennie Fagan for cloning the plasmid expressing SpyCas9. Preliminary experiments were performed during the 2016 CSHL Synthetic Biology summer course. This material is based upon work supported by the Defense Advanced Research Projects Agency (contract HR0011-16-C-01-34, V.N. and C.L.B.), the Office of Naval Research (award N00014-13-1-0074, V.N.), the National Institutes of Health (grant 1R35GM119561-01, C.L.B.), and the National Science Foundation (grant CBET-1403135 to C.L.B.).

## AUTHOR CONTRIBUTIONS

C.L.B. designed the experiments and wrote the paper. V.N. designed the experiments and wrote the paper. C.S.M. designed and performed the experiments and wrote the paper. R.M. designed and performed the experiments and wrote the paper. M.L.L. designed and performed the experiments. T.J. performed the experiments. S.P.C. performed the experiments.

## MATERIALS AND METHODS

### Preparation of TXTL lysates and reactions

The *E. coli* cell-free TXTL lysate was prepared from BL21 Rosetta 2 from Novagen as described previously (Caschera and Noireaux, 2014). The dSpyCas9-loaded TXTL lysate followed the same procedure, only the *E. coli* cells harbored a plasmid constitutively expressing dSpyCas9 from a J23108 promoter. TXTL reactions were composed of 33% volume crude extract and the remaining 67% volume with the following components: energy mix, amino acid mix, cofactors, ions, and DNA. A typical TXTL reaction contains 50 mM HEPES pH 8, 1.5 mM ATP and GTP, 0.9 mM CTP and UTP, 0.2 mg/mL tRNA, 0.26 mM coenzyme A, 0.33 mM NAD, 0.75 mM cAMP, 0.068 mM folinic acid, 1 mM spermidine, 30 mM 3-PGA, 1.5% PEG8000, 30 mM maltodextrin, 3 mM of each of the 20 amino acids, 90 mM K-glutamate, and 4 mM Mg-glutamate. TXTL reactions were conducted in volumes of 5 μl at 29-30°C. When expressing from linear DNA template, 2 μM of annealed oligos containing six copies of the χ-site sequence (Chi6; for details see (Marshall et al., 2017)) was added to the reaction.

### Assembly of expression constructs

Plasmids were constructed using standard techniques. The sequence of each gBlock and plasmid used in this experiment is available in the supporting information (Table S1). The plasmid expressing catalytically active SpyCas9 was generated by amplifying the transcriptional unit expressing SpyCas9 from pCas9 but excluding the CRISPR array from using primers CSMpr1132/1157 and cloning it into the backbone pCSM117. pCas9 was a gift from Luciano Maraffini (Addgene plasmid #42876). gBlocks were ordered from IDT and amplified with CSMpr1105/1106 before being PCR purified. All constructed plasmids were verified by Sanger sequencing of the inserted sequences.

### Fluorescence time-course measurements in TXTL

Fluorescence kinetics measurements were performed principally using the reporter plasmid P70a-deGFP expressing a truncated version of eGFP (deGFP, 25.4 kDa, 1 mg/mL = 39.38 μM) (Shin and Noireaux, 2012). Fluorescence was measured on a Biotek H1m plate reader using a 96-well V-bottom plate (Corning Costar 3357) and Ex 485 nm, Em 528 nm. Time-course measurements were run for at least 16 hours at 29-30°C, with an interval of 3 minutes between reads. Fluorescence intensity measurements were quantified using a linear standard curve, spanning over three orders of magnitude, produced with pure recombinant eGFP (Cell Biolabs Inc., San Diego, CA). Error bars are the standard error of measurement based on at least three replicates run on the same day. GFP production rates were calculated by two-point numerical differentiation and smoothed with a five-point quadratic polynomial. The time to repression was calculated based on the earliest time in which the rates of deGFP production diverge for the targeting sgRNA and the non-targeting sgRNA.

### Amplification of DNA targets in TXTL

TXTL reactions with the SpyCas9 or dSpyCas9 plasmid, sgRNA DNA template, and the P70a-deGFP expression plasmid were incubated at 29°C for three hours, then diluted 200X in water. 1 μl of this dilution was then used as the DNA template in a 50 μl PCR reaction with 55°C annealing temperature, 45 s extension time, and 25 cycles. The PCR reaction was conducted using Taq polymerase and primers RM01s and RM05as to produce a 1074-bp amplicon. PCR products were visualized on a 0.8% agarose gel.

### Fluorescence measurements in *E. coli.*

*E. coli* K-12 MG1655 cells with plasmids expressing dSpyCas9 and deGFP was transformed by electroporation with plasmids encoding the sgRNAs targeting sites (g1-19 targeting) on deGFP, as well as a non-targeting control (g-nt). To measure each strain, three colonies from a freshly streaked plate was inoculated into 2 mL of LB with 34 μg/ml chloramphenicol (Cm), 50 μg/ml ampicillin (Amp), and 50 μg/ml kanamycin (Kan). The strains were cultured for 16-hours at 37° C shaking at 250 rpm. The cultures were then diluted 1:10,000 in LB with Cm, Amp, and Kan to a final volume of 100 μL within a black 96-well assay plate. Each culture was diluted into two wells as technical replicates. Cultures were incubated at 37° C with single orbital shaking at 425 rpm on a 3 mm diameter circle within a BioTek Synergy H1 for 20 hours at which cultures were in stationary phase. Cultures were resuspended with a multichannel pipette and loaded back into the BioTek Synergy H1. Single-point fluorescence at 485/528 nm excitation/emission as well as OD_600_ were measured from each well. Endpoint fluorescence values were background subtracted using the fluorescence from cells lacking deGFP and then normalized by the endpoint, background-subtracted OD_600_ value. Fold repression was then calculated as the normalized fluorescence of the non-targeting sgRNA strain divided by the normalized fluorescence of the targeting sgRNA strain.

### TXTL-based PAM assay

The pET vector expressing FnCpf1 was a kind gift of Benson Hill Biosystems. A PAM library with five randomized nucleotides flanking the spacer sequence was prepared as described previously (Leenay et al., 2016). The crRNA expressing the FnCpf1 crRNA targeting the PAM library was expressed from the gBlock CSM-GB099. A TXTL reaction was assembled as described above except the reaction contained 0.5 mM IPTG and the following DNA components: 0.2 nM P70a-T7RNAP, 2 nM pET-FnCpf1, 2 μM of Chi6 annealed oligos, 0.5 nM of the 5N PAM library, 2 nM of linear DNA expressing the crRNA. The reaction was split into 5 μL reactions as above and incubated at 29°C. Samples were then collected at the specified time by cutting away the cap mat sealing individual reactions and immediately freezing the reaction at −20°C until subsequent use. The adapters were attached by PCR amplification of the TXTL reaction using NEB’s Q5 High-FIdelity DNA Polymerase with RL133 and RL134. The PCR reaction was then purified using Ampure beads before attaching unique Nextera indices to each sample by PCR amplification. After a final PCR purification using Ampure beads, each sample was normalized to 10 nM in a total volume of 20 uL and submitted for next-generation sequencing with 150 single-end reads on an Illumina MiSeq machine. The PAM wheel was generated as described previously (Leenay et al., 2016).

